# Mechanism of substrate recognition and catalysis in a mammalian phosphatidylserine synthase 2

**DOI:** 10.64898/2026.02.06.704487

**Authors:** Lie Wang, Zhen Zhang, Hongyuan Yang, Arthur Laganowsky, Ming Zhou

## Abstract

Mammalian phosphatidylserine synthase-1 and -2 synthesize phosphatidylserine (PS) by replacing the headgroup of either phosphatidylcholine (PC, PTDSS1) or phosphatidylethanolamine (PE, PTDSS2) with a serine. We determined structures of PTDSS2 from *Equus caballus* in complex with either PE or serine substrates to resolutions of 2.8-3.2 Å. The structures define substrate binding sites and reveal that the phosphate group of PE is coordinated by two Ca^2+^. In addition, we found that PTDSS2 has significant phospholipase D (PLD) activity in the absence of serine, which was not reported previously, and that Ca^2+^ is required for the PLD activity. These discoveries enrich our knowledge in the mechanism of mammalian PTDSS.

## Introduction

Phosphatidylserine (PS) is an essential building block of eukaryotic cell membranes and an important signaling molecule. PS accounts for ∼15% of the total phospholipids in cell membranes and is the most abundant negatively-charged phospholipids^1^. PS is located almost exclusively to the intracellular leaflet of the plasma membranes, and exposure of PS to the extracellular leaflet is a signal that triggers important processes such as apoptosis and blood clotting^2–5^. In addition, PS regulates trafficking of cholesterol from plasma membrane to endoplasmic reticulum membrane^6^.

In mammals, PS synthesis is accomplished by two homologous enzymes, phosphatidylserine synthase-1 and -2 (PTDSS1 and PTDSS2), which are ∼32% identical and ∼52 % similar in amino acid sequences^7,8^. PTDSS1 and PTDSS2 are primarily found in the endoplasmic reticulum (ER), and enriched in the mitochondria-associated membrane (MAM) of the ER^9^. Mutations in PTDSS1 are known to cause the Lenz-Majewski syndrome (LMS), symptoms of which include intellectual disability and anomalies in bone development^10^. Recent studies have also validated PTDSS1 as a target for treating cancers^11,12^.

Previous functional studies found that Ca^2+^ is required for PTDSS activities^13–15^. Recently structures of human PTDSS1 and PTDSS2 defined a single Ca^2+^ site that facilitates coordination of serine^16–18^. In the current study, we provide strong evidence that PTDSS has two Ca^2+^ binding sites, and that Ca^2+^ facility coordination of the phosphate group in the PE substrate. The structures also provide a simple mechanistic interpretation of the phospholipase activity in PTDSS2.

## Results

### Functional characterization of PTDSS2

We expressed and purified PTDSS1 from human (hPTDSS1, NCBI reference ID: NM_014754.3) and PTDSS2 from *Equus caballus* (ecPTDSS2, NCBI reference ID: XP_001489141.3). The amino acid sequence of ecPTDSS2 is 90% identical to that of human PTDSS2. Both hsPTDSS1 and ecPTDSS2 elutes as a dimer on a size-exclusion column, and the dimeric assembly is further confirmed by structural studies. Since ecPTDSS2 is more stable after purification, it is the primary focus for structural and functional studies in this report. Addition of cholesteryl hemisuccinate (CHS) during the purification is crucial for the stability of ecPTDSS2, suggesting that ecPTDSS2 may interact with cholesterol (**Extended Data Figs. 1a, b**).

Enzymatic activity of purified ecPTDSS2 is demonstrated by following the conversion of an NBD-labeled PE to NBD-PS on a thin-layer chromatography (TLC, **Fig. 1a, b**). Ca^2+^ is required for the reaction, and interestingly, in the absence of serine and in the presence of Ca^2+^, ecPTDSS2 exhibits phospholipase D (PLD) activity that leads to the production of PA (**Fig. 1c**). ecPTDSS2 shows high specificity for PE, as free ethanolamine (2 mM) inhibits ecPTDSS2 activity while free choline (2 mM) does not (**Fig. 1d**). ecPTDSS2 accepts threonine as a substrate but the reaction seems slower, but not tyrosine or cysteine (**Fig. 1e**). Ca^2+^ is required for enzymatic activity, and neither Mg^2+^, Mn^2+^ or Zn^2+^ can replace the role of Ca^2+^ in catalysis (**Fig. 1f**).

**Figure 1.**
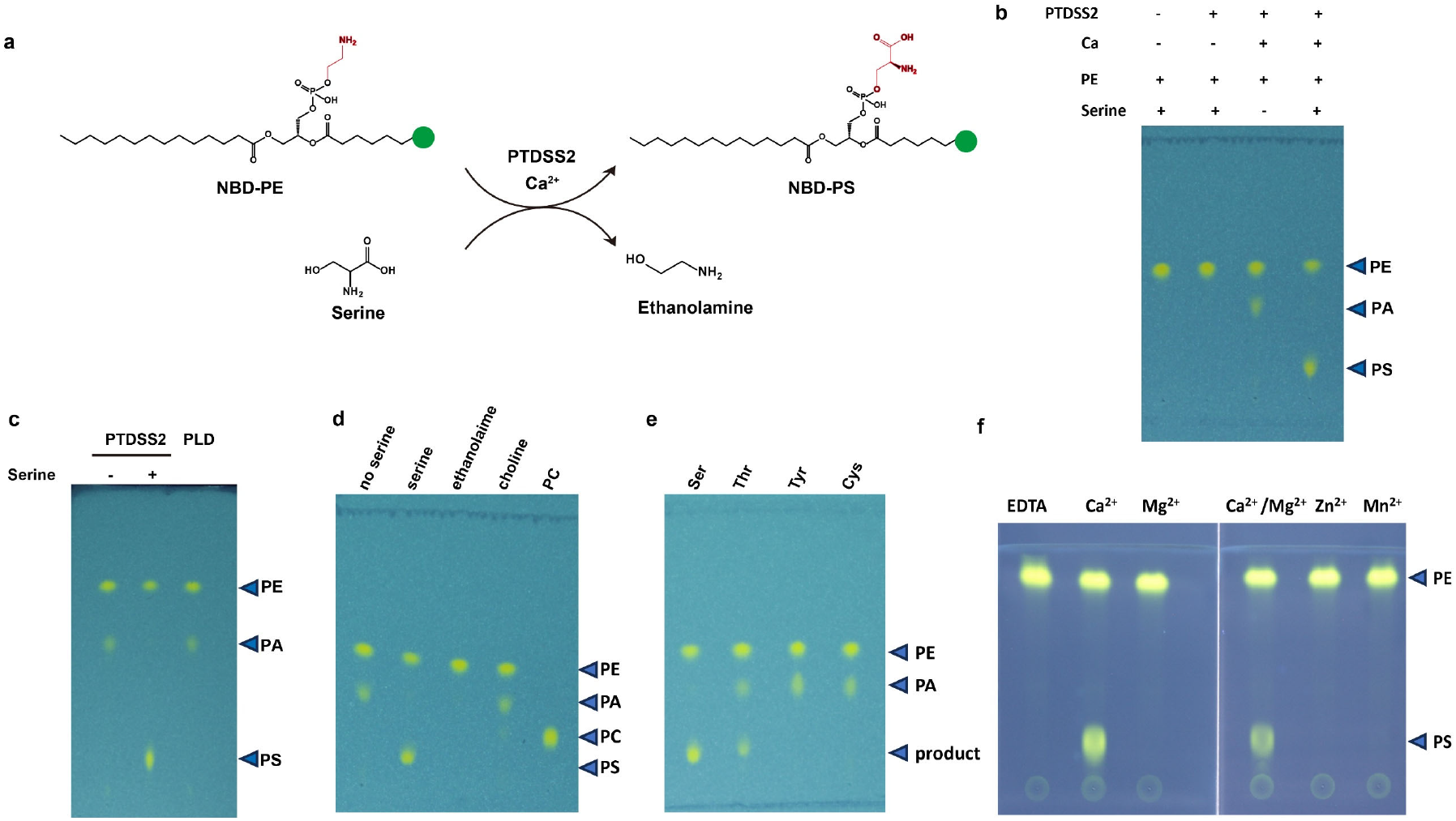
Function of ecPTDSS2. **a**. Reaction catalyzed by ecPTDSS2. **b**. Production of PS in the presence of Ca, PE and serine. **c**. PE-PLD activity of ecPTDSS2. **d-e**. Substrate selectivity of ecPTDSS2 to PE and serine. In first 4 lanes of 1d, PE was used as headgroup acceptor, and PC and serine were used in the last lane. In 1e, PE was used as headgroup acceptor. **f**. Cation selectivity of ecPTDSS2.

### Overall structure of ecPTDSS2

We determined the structures of ecPTDSS2 using single-particle cryo-electron microscopy (cryo-EM) under three different conditions, in the presence of Ca^2+^ and serine at ∼2.9 Å resolution, of Ca^2+^ and POPE at ∼3.2 Å, and of Ca^2+^ and NBD-PA at ∼2.8 Å (**Extended Data Fig. 2-4, table 1**).

The three density maps are of sufficient quality to allow *de novo* model building of most of ecPTDSS2 including all 10 transmembrane helices (**Extended Data Fig. 5a**), except for the first 62 residues (1-62), the last 75 residues (433 to 498), and the loop between TM6 and TM7 (281 to 297), which are not resolved. Since the three structures are almost identical with overall root-mean-squared deviations (RMSD) of less than 0.6 Å, we will use the structure of ecPTDSS2 in the presence of Ca^2+^ and serine for the purpose of illustrations in all the figures except for Extended Data Figure 9, which shows ecPTDSS2 in complex with NBD-PA.

ecPTDSS2 forms a homodimer and each protomer has 10 transmembrane helices. Based on the “positive-inside” rule^19^, both the N- and C-termini are on the cytosolic side (**Extended Data Fig. 5b**). This assignment is further supported by the location of the two Ca^2+^ binding sites and a structurally resolved disulfide bridge between Cys172 and Cys239, both of which are exposed to the lumen side of the ER. The 10 helices segregate into two well-defined domains: TM1-3 and TM9 form a panel-like structure that serves as the dimer interface, and hence the dimerization domain; TM4-8 fold into a cone-like structure that encloses the substrate binding sites and the catalytic residues, and hence the catalytic domain (**Fig. 2c, d**). TM10 seems suspended in the membrane and interacts only with TM5 and there is a partially resolved cholesterol molecule at the interface.

**Figure 2.**
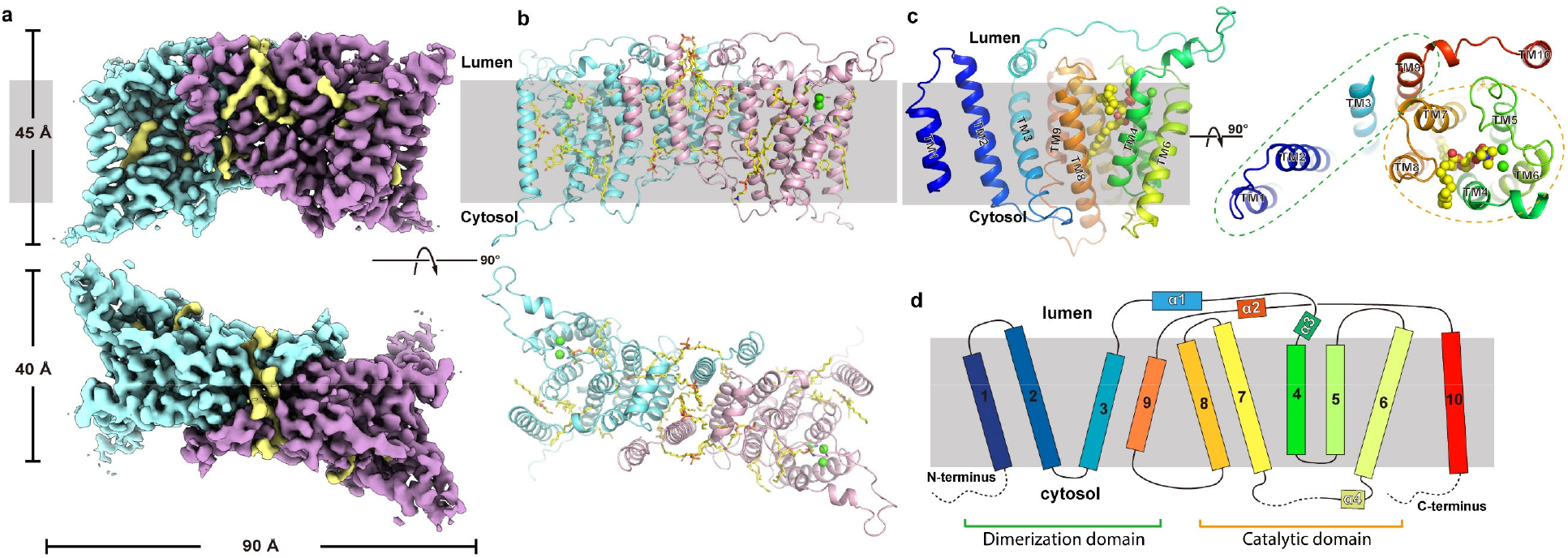
Overall structure of ecPTDSS2. **a**. The density map of dimeric ecPTDSS2. The two monomers are colored in cyan and pink, and densities of lipids and substrates are colored in yellow. **b**. Structure model of the dimeric ecPTDSS2. Two monomers are shown as cartoon in cyan and pink, and the substrates and lipids are shown as yellow sticks. Two Ca^2+^ in each monomer are shown as green spheres. **c**. Structure of the monomer in two orientations. **d**. Topology plot of an ecPTDSS2 monomer.

### The catalytic domain: two Ca^2+^ binding sites and lateral entry of PE

The open end of the cone-shaped catalytic domain faces the luminal side of the ER (**Fig. 2c**). Conserved negatively charged residues, Glu223, Glu226 from TM5, Glu238, Asp242 and Asp247 from TM6, cluster at the open end. Two non-protein densities are present in the vicinity of the negatively charged residues, and we assign each with a Ca^2+^, Ca1 and Ca2 (**Fig. 3a-b**). Ca1 is coordinated by the carboxylates of Glu223 and Glu 226, while Ca2 is coordinated by the carboxylates of Asp242 and Asp247. The carboxylate of Glu238 seems to bridge both Ca^2+^. The relative intensity of the two Ca^2+^ densities differs with Ca1 at 18σ and Ca2 at 14σ, comparable to that of protein backbone in the vicinity. We made alanine mutations, one at a time, to residues Glu223, Glu238 and Asp247 in ecPTDSS2, and found that all three mutant proteins lose the enzymatic activity (**Fig. 3c**). A similar mutational study on PTDSS1 from Chinese hamster also show that all five negatively charge residues are crucial for the enzymatic activity^14^.

**Figure 3.**
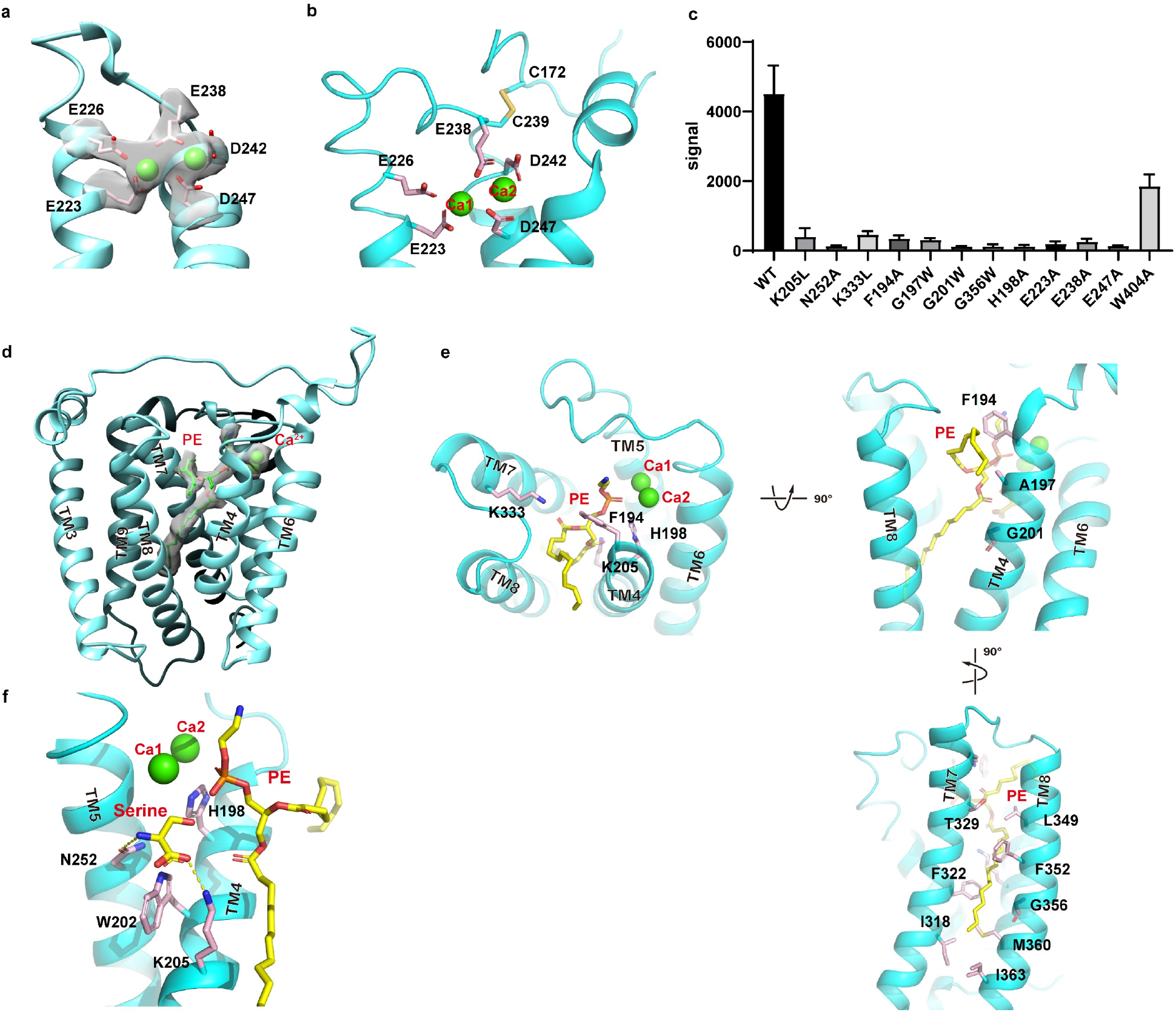
Ca^2+^ and substrate binding sites. **a. & b**. The Ca^2+^ binding sites. Density map of side chains and Ca^2+^ are shown in gray, Ca^2+^ as green spheres, and side chains coordinating Ca^2+^ are shown as pink sticks. A disulfide bridge between Cys 172 and 239 is shown in yellow sticks. **c**. Enzymatic activity. ^14^C-signal of the PS product is plotted for the wild type and mutants. Each bar is the average of three independent measurements, and error bars are standard errors of the mean (s.e.m.). **d & e**. PE binding site. Density of PE is shown as gray transparent surface, PE in yellow sticks, and Ca^2+^ as green spheres. PE and Ca^2+^ are shown in three views. **f**. The complete active site assembled from different structures. The relative positions of serine, PE, Ca^2+^ and His198 in ecPTDSS2.

The cone-shaped catalytic domain has a lateral opening between TM4 and 8 that provides access to the lipid bilayer (**Figure 3d**). Assignment a serine and a partial POPE to the non-protein densities in the catalytic domain is facilitated by comparison of density maps of samples with or without a substrate (**Extended Data Fig. 6**).

The bound POPE is well resolved except for its 2-oleoyl side chain (**Fig. 3d, Extended Data Fig. 6**). The phosphate group seems stabilized by the two Ca^2+^, while the amine group by the aromatic ring of F194 through cation-π interactions. The carbonyl oxygen atoms from the two acyl chains are each coordinated by the side chains of Lys205 and Lys333 (**Fig. 3e**). The 1-acyl chain resides inside of the cone and is sheathed by a hydrophobic pocket formed by residues mainly from TM7 and 9, while the 2-acyl chain is partially resolved and extends through the lateral opening to the core of the membrane. The importance of residues forming the PE binding site are demonstrated by functional analysis: Single mutation to residues that coordinate the carbonyl oxygens, K205A or K333A, bulky side chains that could impede lipid entry into the cone, G197W and G201W, and a point mutation that reduces the hydrophobic binding pocket for the 1 acyl chain, G356W, all significantly reduce the enzymatic activity (**Fig. 3c**).

The serine is stabilized by the side chains of Lys205 and Asn252. The hydroxyl oxygen of the serine is 2.7 Å from the highly conserved His198 and 4.2 Å from the phosphate atom of the POPE (**Fig. 3f**). This arrangement suggests that His198 is a catalytic residue that activates the hydroxyl oxygen of serine, and the two Ca^2+^ stabilize the binding of the phosphate group on a PE substrate. Mutations to His198, Lys205, and Asn252 significantly reduce the enzymatic activity.

### Dimerization interface and bound lipids

The dimerization interface is formed between the two dimerization domains, in which TM1-2 of one protomer make contacts with TM3 and 9 of the neighboring protomer. The interactions at the dimer interface are mainly of hydrophobic side chains (**Fig. 4a**). In addition, there is an enclosed narrow space between the two promoters that is filled with phospholipid-like densities, into which we built six partial lipid molecules (**Fig. 4b**).

**Figure 4.**
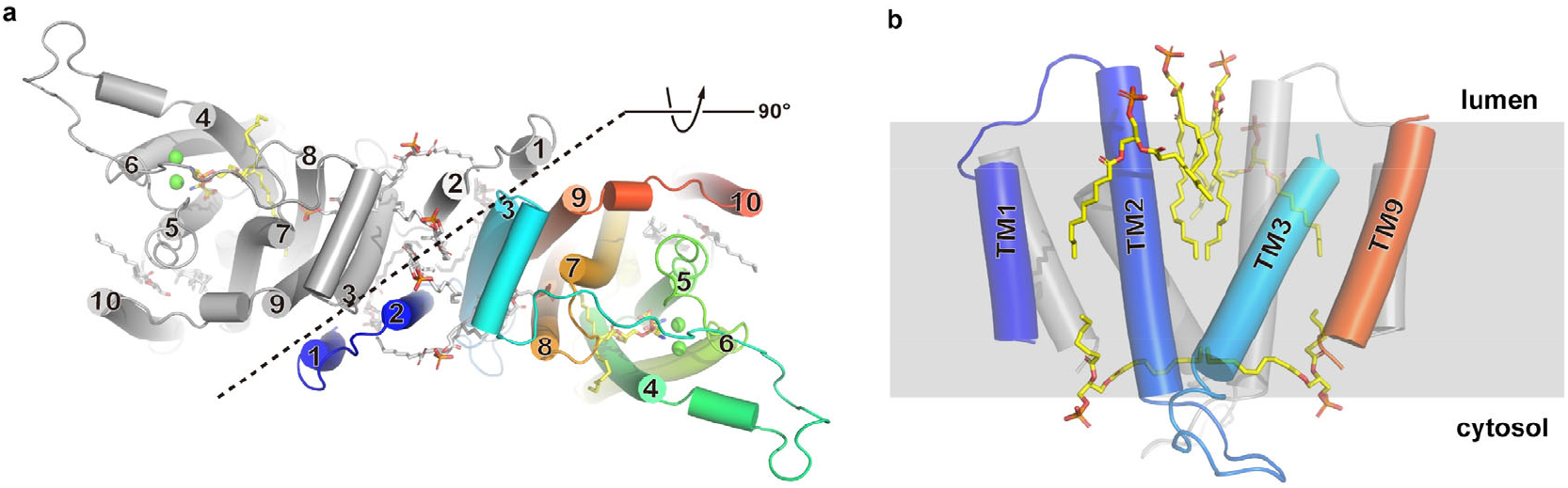
Dimerization interface and bound lipids. **a**. ecPTDSS2 is shown as cartoon, with one protomer colored in gray and another in rainbow. Each TM is labeled on the end, and lipids are shown as yellow sticks. **b**. Lipids resolved at the dimer interface. The dimerization domains are shown in gray cartoon for one subunit and rainbow cartoon for another. Lipids are shown as yellow sticks.

## Discussion

In summary, we expressed and purified ecPTDSS2 and demonstrated its enzymatic activity. We determined structures of ecPTDSS2 and defined binding sites for the two substrates, PE and serine. The structures also identified two Ca^2+^ binding sites. These observations provide a foundation for envisioning a mechanism of PS synthesis in eukaryotic cells: A PTDSS engages two Ca^2+^ with the highly conserved acidic residues located on the luminal side of ER; Serine likely enters its binding site from the intracellular side, and its hydroxyl group is activated by the highly conserved catalytic His198; A PE located to the luminal leaflet of the ER membrane diffuses into its binding site through a lateral opening between TM4 and TM8, and its phosphate group is stabilized by the two Ca^2+^ and its acyl chain accommodated by a deep hydrophobic pocket formed by TM7 and TM8. Nucleophilic attack of the activated hydroxyl oxygen of serine on the phosphate of PE leads to production of PS and detachment of ethanolamine (**Fig. 5a**). In this model, the two Ca^2+^ enhance the binding of the phosphate group and stabilize the reaction intermediate during the nucleophilic attack.

**Figure 5.**
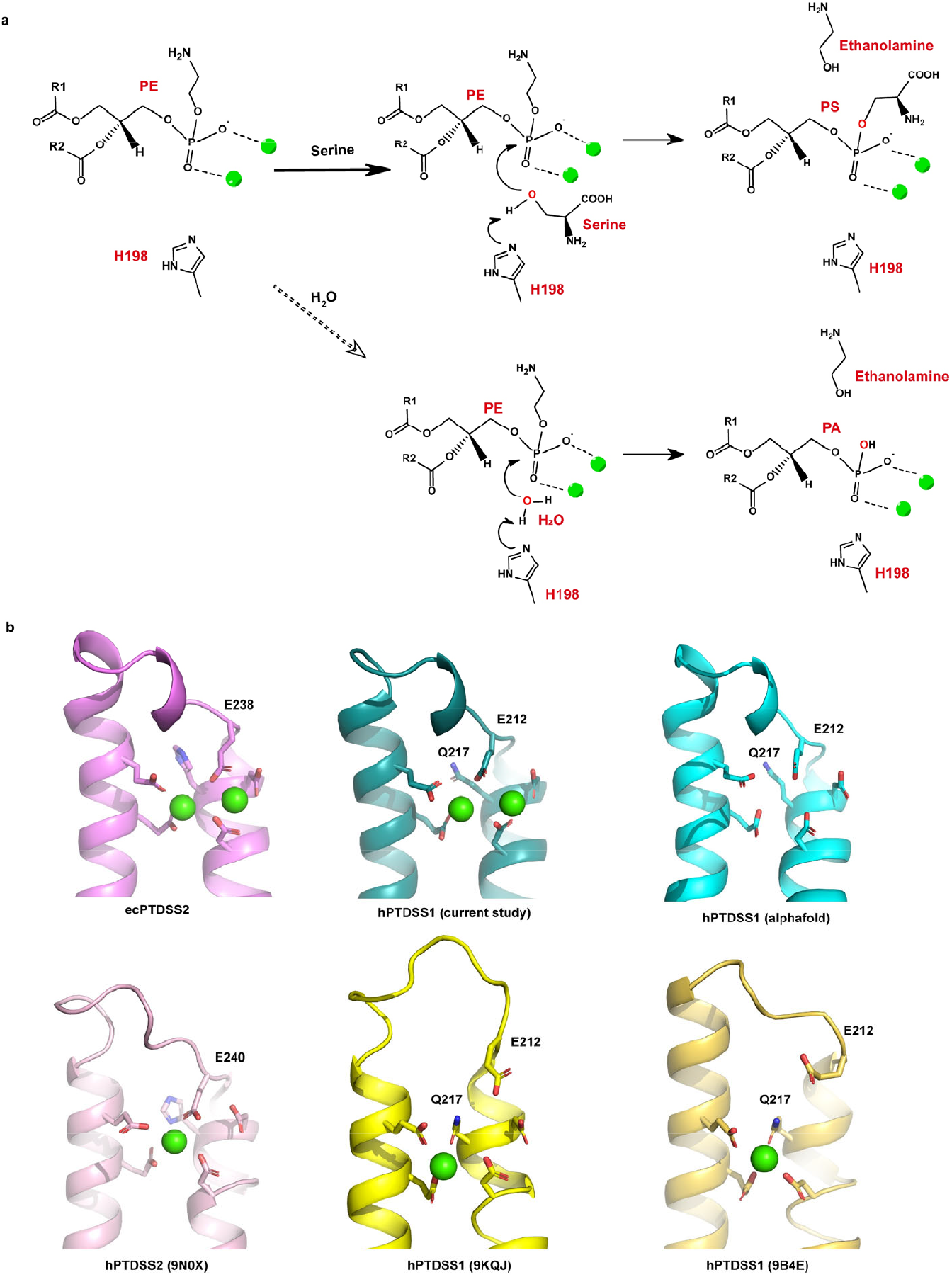
Proposed mechanism of reaction. **a**. Proposed catalytic mechanism of PTDSSs. Top panel, Proposed coordination of PE by two Ca^2+^ (dashed lines) and activation of serine hydroxyl by H198 (arrows); Bottom panel, In the absence of serine, a water molecule replaces serine to promote hydrolysis. **b**. Review of Ca^2+^ binding sites in known PTDSS structures. Ca^2+^ binding sites in the structures of ecPTDSS2 (purple), hPTDSS1 (teal), AF3 predicted hPTDSS1 (cyan), PTDSS2 (pink, PDB ID: 9N0X), PTDSS1 (yellow, PDB ID: 9KQJ), and PTDSS1 (gold, PDB ID: 9B4E). Ca^2+^ ions are shown as green spheres, and residues that are involved in Ca^2+^ binding are shown as sticks.

We are aware that in previous publications of human PTDSS1 & 2, the proposed catalytic mechanism also identifies the equivalent histidine as a key residue for activating the hydroxyl group, however, these structures identified a single Ca^2+^. Due to different placement of the serine substrate, the Ca^2+^ was assigned to coordinate the carboxylate group of serine^16,18^. A comparison of the two Ca^2+^ binding sites in the current structures to the single Ca^2+^ site identified in previously reported PTDSS structures are shown in **Fig. 5b**. The positions of at least 4 out of 5 highly conserved acidic residues are identical in these structures, with the 5^th^ acidic residue, Glu238 (PTDSS2 numbering) also in the same position as that of human PTDSS2 (PDB ID 9N0X) but in a different position to that of the other two structures (PDB ID 9KQJ and 9B4E). In our opinion, binding of two Ca^2+^ is more compatible with the cluster of five highly conserved carboxylates arranged in proximity and thus is a general feature of this family of enzymes. Indeed, when we expressed and purified human PTDSS1 and determined its structure in the presence of 2mM Ca^2+^ to ∼3.1 Å resolution (**Extended Data Fig. 7**), we found two Ca^2+^ identically positioned to these of ecPTDSS2 (**Fig. 5b and Extended Data Fig. 8**). Interestingly, AlphaFold3 also predicts similar arrangement of all 5 acidic residues as reported in this study and in ref 14.

We are intrigued by the observation that hydrolysis of PE occurs in the absence of serine (**Figure 1b**), and we explored this further with the structural study. We determined the structure of ecPTDSS2 after its incubation with NBD-PE in the absence of serine to a resolution of 2.8 Å (**Extended Data Fig. 4-5**). Although the structure did not capture a reaction intermediate, it clearly shows the presence of NBD-PA, which is the hydrolysis product of NBD-PE. The unlabeled 1-acyl chain is buried in the hydrophobic pocket while the NBD-labeled 2-acyl chain straddles the entrance between TM4 and TM8, consistent with their assignments in the structure of ecPTDSS2-POPE. In addition, NBD-PA is captured in a position a few angstroms away from the Ca^2+^ sites so that its phosphate is no longer coordinated, indicating that we may have captured an intermediate state in the process of product dissociation (**Extended Data Fig. 6, 9a**). The structure also shows that TM4 is shifted by a few tenth of an Angstrom away from TM8 when compared to that of ecPTDSS2-POPE (**Extended Data Fig. 9b, c**), suggesting that a mobile TM4 is a required element for substrate entry or product egress.

## Methods

### Cloning, expression and purification of *Equus caballus* PTDSS2

Condon-optimized ecPTDSS2 gene (NCBI reference ID: XP_001489141.3) was cloned into the pFastBac-Dual expression vector for baculovirus production^20^. The P3 virus was used to infect High Five™ (Trichoplusia ni) cells at a density of 3×10^6^ cells/ml and the cells were harvested after 60 hrs. Cell membranes were prepared by a hypotonic/hypertonic wash protocol as previously described^21^. Briefly, cells were lysed in a hypotonic buffer containing 10 mM 4-(2-hydroxyethyl)-1-piperazineethanesulfonic acid (HEPES, pH 7.5), 10 mM NaCl, and 2 mM β-mercaptoethanol (BME), and 25 μg/ml DNase I. After ultracentrifugation at 55,000 × g for 10 min, the pelleted cell membranes were resuspended in a hypertonic buffer containing 25 mM HEPES pH 7.5, 1 M NaCl, and 25 μg/ml DNase I, and were centrifuged again at 55,000 × g for 20 min. The pelleted cell membranes were resuspended in 20 mM HEPES pH 7.5, 150 mM NaCl, 20 % glycerol and flash frozen in liquid nitrogen.

Purified membranes were thawed and extracted in 20 mM HEPES pH 7.5, 150 mM NaCl, 4 mM CaCl_2_, 2 mM beta-mercaptoethanol, 1 tablet of protease inhibitors (Roche) and then solubilized with 1.5% (w/v) lauryl maltose neopentyl glycol (LMNG, Anatrace) at 4 °C for 2 h. After solubilization, cell debris was removed by centrifugation in 55,000 × g, 45 min, 4 °C, ecPTDSS2 was purified from the supernatant with a cobalt-based affinity resin (Talon, Clontech). The C-terminal His6-tag was cleaved with tobacco etch virus protease at room temperature for 30 min. The protein was concentrated to ∼5mg/ml (Amicon 100 kDa cut-off, Millipore), and loaded onto a size-exclusion column (SRT-3C SEC-300, Sepax Technologies) equilibrated with 20 mM HEPES, 150 mM NaCl, 4 mM CaCl_2_ and 0.1% (w/v) LMNG, 0.01 (w/v) cholesteryl hemisuccinate (CHS, avanti). For samples used in cryo-EM, the size-exclusion column was equilibrated with 20 mM HEPES, 150 mM NaCl, 2 mM beta-mercaptoethanol, 4 mM CaCl_2_ and 0.02% (w/v) GDN. Human PTDSS1 was expressed and purified with a similar protocol. For the ecPTDSS2-Ca^2+^-Serine sample, 5 mM Serine was added before grid preparation. For the ecPTDSS2-Ca^2+^-POPE sample, 0.5 mM POPE were added before grid preparation. For ecPTDSS2-Ca^2+^-NBD-PA sample, 0.1 mM NBD-PE was added before loading onto the size-exclusion column.

ecPTDSS2 mutants were generated using the QuikChange method (Stratagene) and the entire cDNA was sequenced to verify the mutation. Primer sequences are provided in the Source Data file. Mutants were expressed and purified following the same protocol as wild type.

### Cryo-EM sample preparation and data collection

Cryo-grids were prepared on a Thermo Fisher Vitrobot Mark IV. Quantifoil R1.2/1.3 Cu grids were glow-discharged using the Pelco Easyglow. Concentrated ecPTDSS2 (3.5 μl) was applied to glow-discharged grids. After blotting with filter paper (Ted Pella) for 3.5-4.5 s, the grids were plunged into liquid-nitrogen-cooled liquid ethane. For cryo-EM data collection, movie stacks were collected using EPU (Thermo Fisher Scientific) on a Titan Krios at 300 kV with a Quantum energy filter (Gatan), at a nominal magnification of ×105,000 and with defocus values of -2.2 to -0.8 μm. A K3 Summit direct electron detector (Gatan) was paired with the microscope. Each stack was collected in the super-resolution mode with an exposing time of 0.175 s per frame for a total of 50 frames. The dose was about 50 e^-^ per Å^2^ for each stack. The stacks were motion-corrected with Relion 4 and binned (2 × 2) ^22^. Dose weighting was performed during motion correction, and the defocus values were estimated with Gctf^23^.

### Cryo-EM data processing

For ecPTDSS2-Ca^2+^-Serine data set, a total of 8,734,375 particles were automatically picked in RELION 4.0 from 8944 images^24^, and imported into cryoSPARC^25^. Out of 200 two-dimensional (2D) classes, 35 classes (containing 1,325,838 particles) were selected for *ab initio* three-dimensional (3D) reconstruction, which produced one good class with recognizable structural features and two bad classes that did not have structural features. Both the good and bad classes were used as references in the heterogeneous refinement (cryoSPARC) and yielded a good class at 4.00 Å from 347,969 particles. After Bayesian polishing of the particles in RELION, nonuniform refinement (cryoSPARC) was performed with C2 symmetry, an adaptive solvent mask, and CTF refinement to produce a map with an overall resolution of 2.9 Å.

For ecPTDSS2-Ca^2+^-POPE/Serine data set, a total of 6,266,551 particles were automatically picked in RELION 4.0 from 6620 images^24^, and imported into cryoSPARC^25^. Out of 200 two-dimensional (2D) classes, 25 classes (containing 1,137,978 particles) were selected for ab initio three-dimensional (3D) reconstruction, which produced one good class with recognizable structural features and two bad classes that did not have structural features. Both the good and bad classes were used as references in the heterogeneous refinement (cryoSPARC) and yielded a good class at 4.26 Å from 279,188 particles. After Bayesian polishing in RELION, nonuniform refinement (cryoSPARC) was performed with C2 symmetry and an adaptive solvent mask, and CTF refinement yielded a map with an overall resolution of 3.2 Å.

For ecPTDSS2-Ca^2+^-NBD-PA data set, a total of 6,602,250 particles were automatically picked in RELION 4.0 from 8974 images^24^, and imported into cryoSPARC^25^. Out of 200 two-dimensional (2D) classes, 30 classes (containing 1,518,283 particles) were selected for ab initio three-dimensional (3D) reconstruction, which produced one good class with recognizable structural features and two bad classes that did not have structural features. Both the good and bad classes were used as references in the heterogeneous refinement (cryoSPARC) and yielded a good class at 3.96 Å from 482,496 particles. After Bayesian polishing in RELION, nonuniform refinement (cryoSPARC) was performed with C2 symmetry, an adaptive solvent mask, and CTF refinement to yield a map with an overall resolution of 2.77 Å.

For hPTDSS1 data set, a total of 2,097,926 particles were automatically picked in RELION 4.0 from 12080 images^24^, and imported into cryoSPARC^25^. Out of 200 two-dimensional (2D) classes, 25 classes (containing 1,034,264 particles) were selected for ab initio three-dimensional (3D) reconstruction, which produced one good class with recognizable structural features and two bad classes that did not have structural features. Both the good and bad classes were used as references in the heterogeneous refinement (cryoSPARC) and yielded a good class at 4.17 Å from 340,120 particles. After Bayesian polishing in RELION, nonuniform refinement (cryoSPARC) was performed with C2 symmetry, an adaptive solvent mask, and CTF refinement to yield a map with an overall resolution of 3.13 Å.

Resolutions were estimated using the gold-standard Fourier shell correlation with a 0.143 cut-off^26^ and high-resolution noise substitution^27^. Local resolution was estimated using ResMap^28^.

### Model building and refinement

The structural models of ecPTDSS2 were built based on prediction model of AlphaFold 2^29^, and sidechains were then adjusted based on the map. Model building was conducted in Coot^30^. Structural refinements were carried out in PHENIX in real space with secondary structure and geometry restraints^31^. The EMRinger Score was calculated as described^32^.

### NBD-PE Enzymatic assay

ecPTDSS2 activity was measured using a fluorescence-based assay. All reaction assays were done in a buffer with 20 mM HEPES, pH 7.5, 150 mM NaCl, 0.02% GDN, 4 mM CaCl_2_. In a single point activity assay, final concentrations of serine and NBD-PE were 10 mM and 0.1 mM, respectively. Reactions were initiated with the addition of 10μg purified ecPTDSS2 protein. After reaction at 37 °C for 30 mins, reactions were stopped by adding extraction solution (methanol: chloroform: water 25:65:4). After vortex and centrifugation, the organic phase was taken for TLC.

### ^14^C-Serine Enzymatic assay

Enzymatic activity was also measured using ^14^C-labeled serine (American Radiolabelled Chemicals, Inc) as a substrate. The assay buffer contains 20 mM HEPES, pH 7.5, 150 mM NaCl, 0.02% GDN, and 4 mM CaCl_2_. Final concentrations of ^14^C-labeled serine and POPE were 0.1 mM and 1 mM, respectively. Reactions were initiated with the addition of 10μg purified ecPTDSS2 protein. After reaction at 37 °C for 30 mins, reactions were stopped by adding extraction solution (methanol: chloroform: water 25:65:4). After vortex and centrifugation, the organic phase was taken, and the radioactivity in the organic phase was determined by liquid scintillation counting.

## Data availability

The atomic coordinates of ecPTDSS2 in complex with Ca^2+^ and serine have been deposited in the PDB (http://www.rcsb.org) under the accession code 10BY. The atomic coordinates of ecPTDSS2 in complex with Ca^2+^ and POPE have been deposited in the PDB under the accession code 10BW. The atomic coordinates of ecPTDSS2 in complex with Ca^2+^ and NBD-PA have been deposited in the PDB under the accession code 10BX. The atomic coordinates of hPTDSS1 in complex with Ca^2+^ have been deposited in the PDB under the accession code 10BZ. Their corresponding electron microscopy maps have been deposited in the Electron Microscopy Data Bank (https://www.ebi.ac.uk/pdbe/emdb/) under the accession codes EMD-75056, EMD-75054, EMD-75055 and EMD-75057, respectively.

## Acknowledgements

This work was supported by grants from NIH (DK122784 to M.Z., GM145416 to A.L. and M.Z., and GM151548 to M.Q. and M.Z.). We acknowledge the cryo-EM cores in Baylor College of Medicine (CPRIT Core Facility Award RP190602) for the support in grid preparation and screening. We thank Dr. Gaya P. Yadav for cryo-EM data collection at the Laboratory for Biomolecular Structure and Dynamics (LBSD) of Texas A&M University. The LBSD is supported, in part, by the Department of Biochemistry & Biophysics, AgriLife, and Texas A&M University. We are grateful to Pacific Northwest Center for Cryo-EM supported by NIH grant U24GM129547. We thank the Laboratory for BioMolecular Structure (LBMS) supported by the DOE Office of Biological and Environmental Research (KP1607011) for support in data collection.

## Author contributions

M.Z. and L.W. conceived and designed the project. L.W. and Z.Z. performed biochemistry experiments, L.W. performed cryo-EM data collection and processing, and model building; M.Z., L.W., Z.Z., A.L., and HY.Y. analyzed the data and wrote the manuscript.

## Extended Data Figures

**Extended Data Figure 1.**
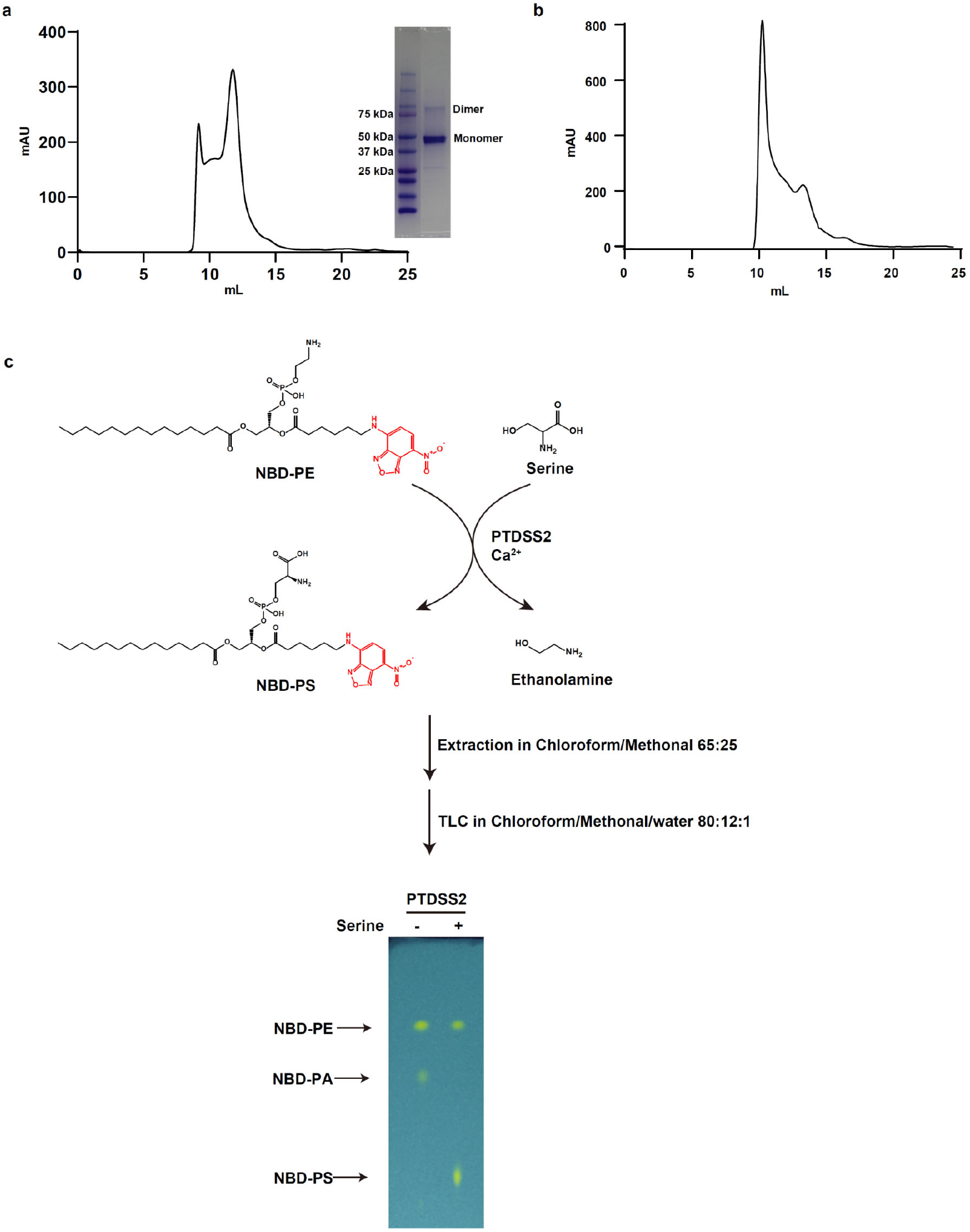
Purification and function characterization of ecPTDSS2. a-b. ecPTDSS2 purification profile with and without presence of CHS. **c**. Function assay with NBD-PE.

**Extended Data Figure 2.**
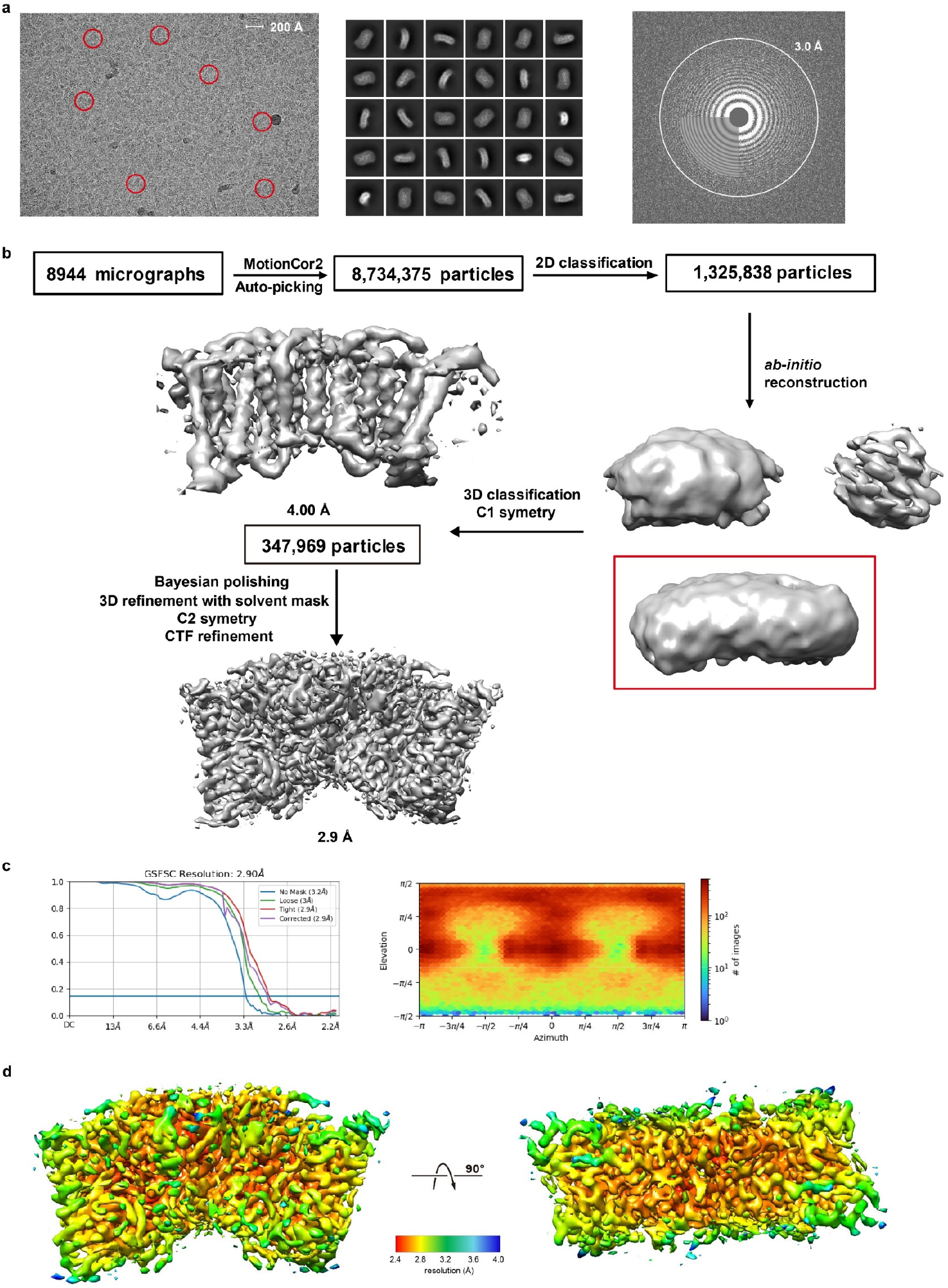
Data processing and map resolution of ecPTDSS2-Serine data set. **a**. A representative micrograph of ecPTDSS2 (left), its Fourier transform (right) and representative 2D class averages (middle). Representative particles are highlighted in red circles. **b**. A flowchart for data processing and the final map. **c**. The gold-standard Fourier shell correlation curve for the final map. **d**. Local resolution map shown in two orientations.

**Extended Data Figure 3.**
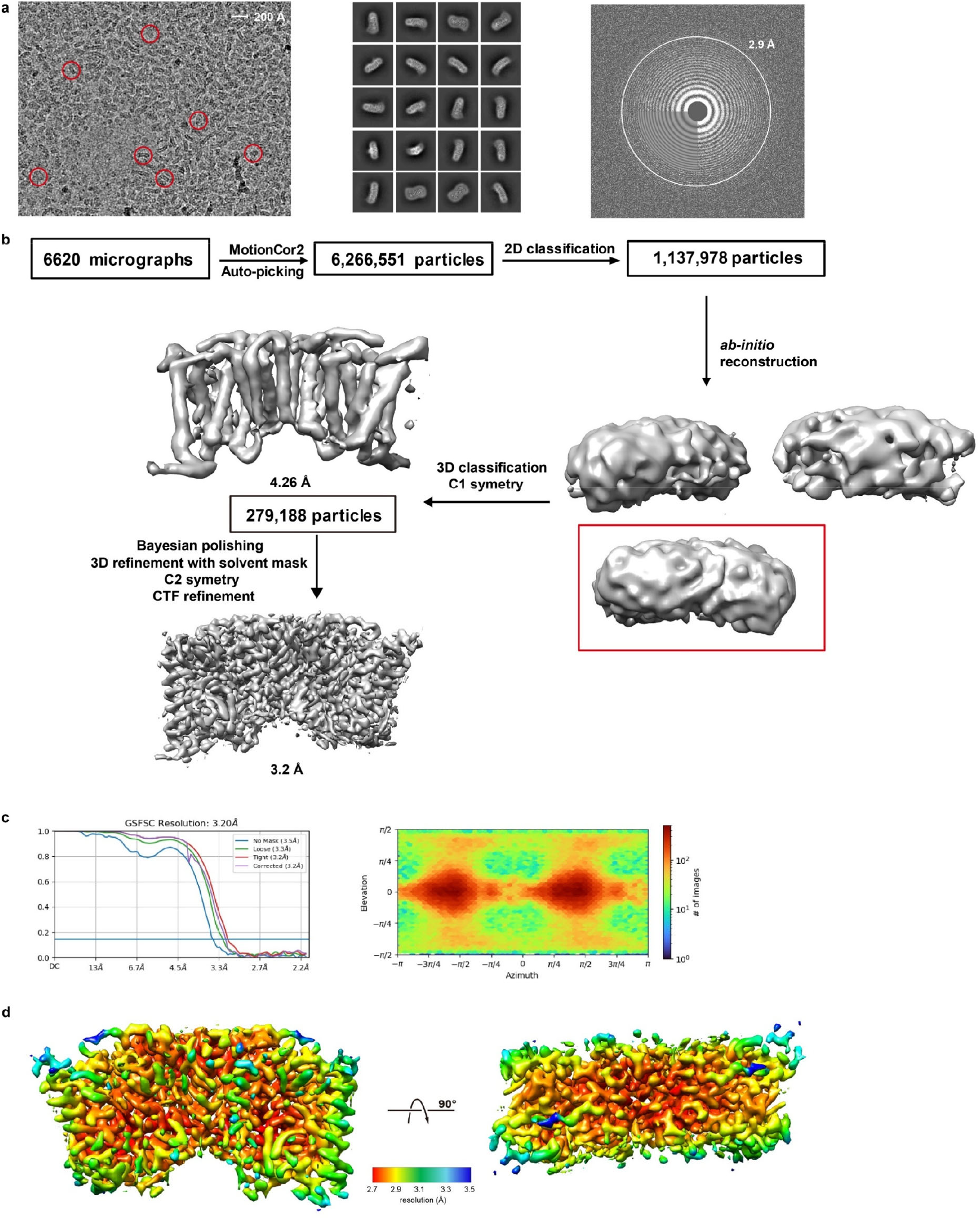
Data processing and map resolution of ecPTDSS2-POPE data set. **a**. A representative micrograph of ecPTDSS2 (left), its Fourier transform (right) and representative 2D class averages (middle). Representative particles are highlighted in red circles. **b**. A flowchart for data processing and the final map. **c**. The gold-standard Fourier shell correlation curve for the final map. **d**. Local resolution map shown in two orientations.

**Extended Data Figure 4.**
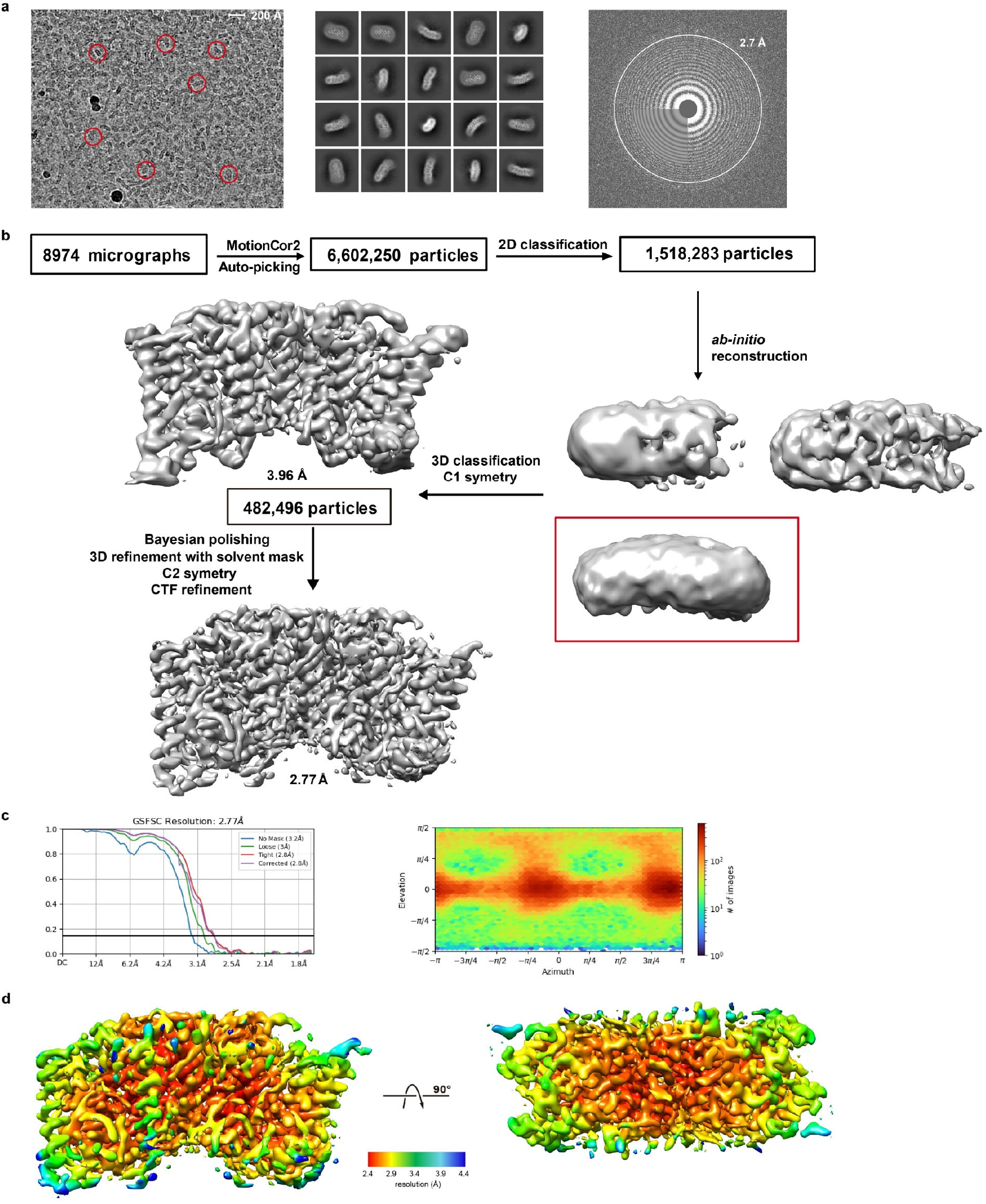
Data processing and map resolution of ecPTDSS2-NBD-PA data set. **a**. A representative micrograph of ecPTDSS2 (left), its Fourier transform (right) and representative 2D class averages (middle). Representative particles are highlighted in red circles. **b**. A flowchart for data processing and the final map. **c**. The gold-standard Fourier shell correlation curve for the final map. **d**. Local resolution map shown in two orientations.

**Extended Data Figure 5.**
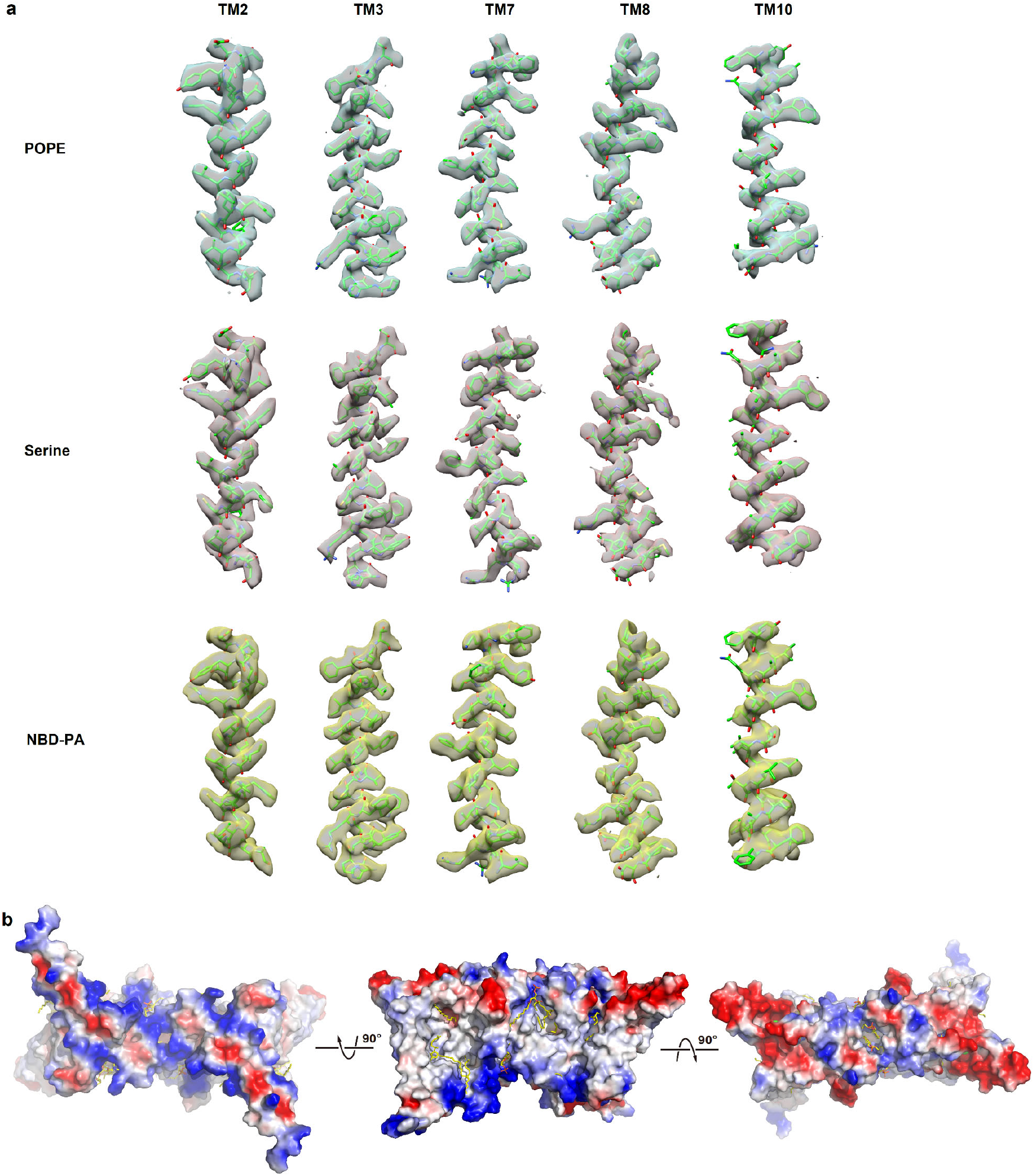
Density maps and structural model of ecPTDSS2. **a**. Individual secondary structures of ecPTDSS2 are shown as green sticks, contoured in their density. Density of the PE-Serine data set is shown as gray surface, density of Serine data set is shown in pink surface, and that of NBD-PE data set is shown as yellow surface. **b**. Electrostatic surface representations of ecPTDSS2 dimer in three orientations. The electrostatic potential is calculated using the APBS plugin from Pymol^33^.

**Extended Data Figure 6.**
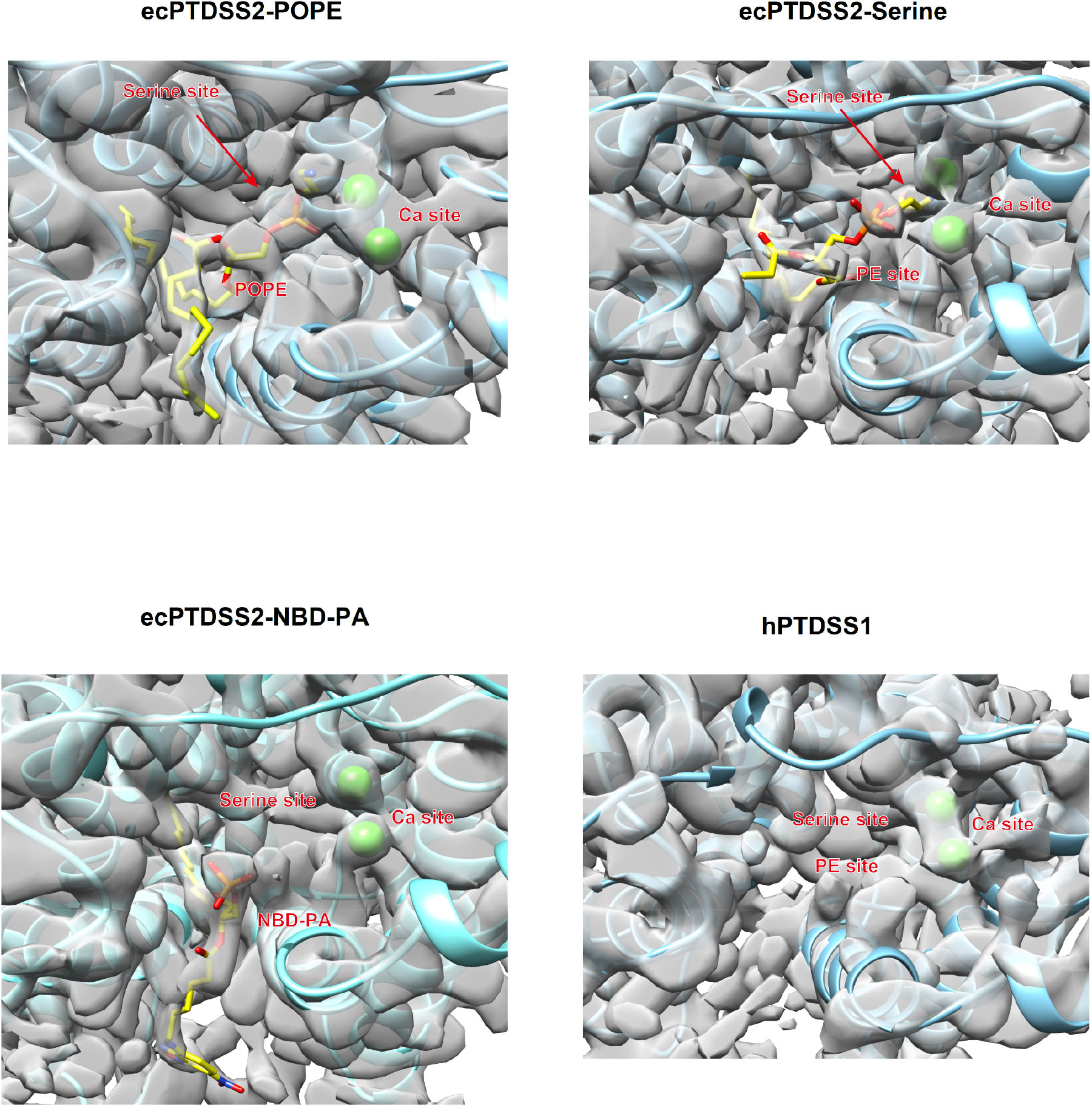
Density comparison of maps of PTDSS. Density is shown as gray transparent surface, protein is shown in cyan cartoon, Ca^2+^ is shown in green sphere, and substrates are shown as yellow sticks. Serine site is occupied in ecPTDSS2-serine map, while it is empty in all other maps. POPE density can be clearly identified in the proposed PE binding site in ecPTDSS2-POPE map, but PE site is empty in NBD-PA and hPTDSS1 map. Two Ca^2+^ sites can be clearly identified in all maps.

**Extended Data Figure 7.**
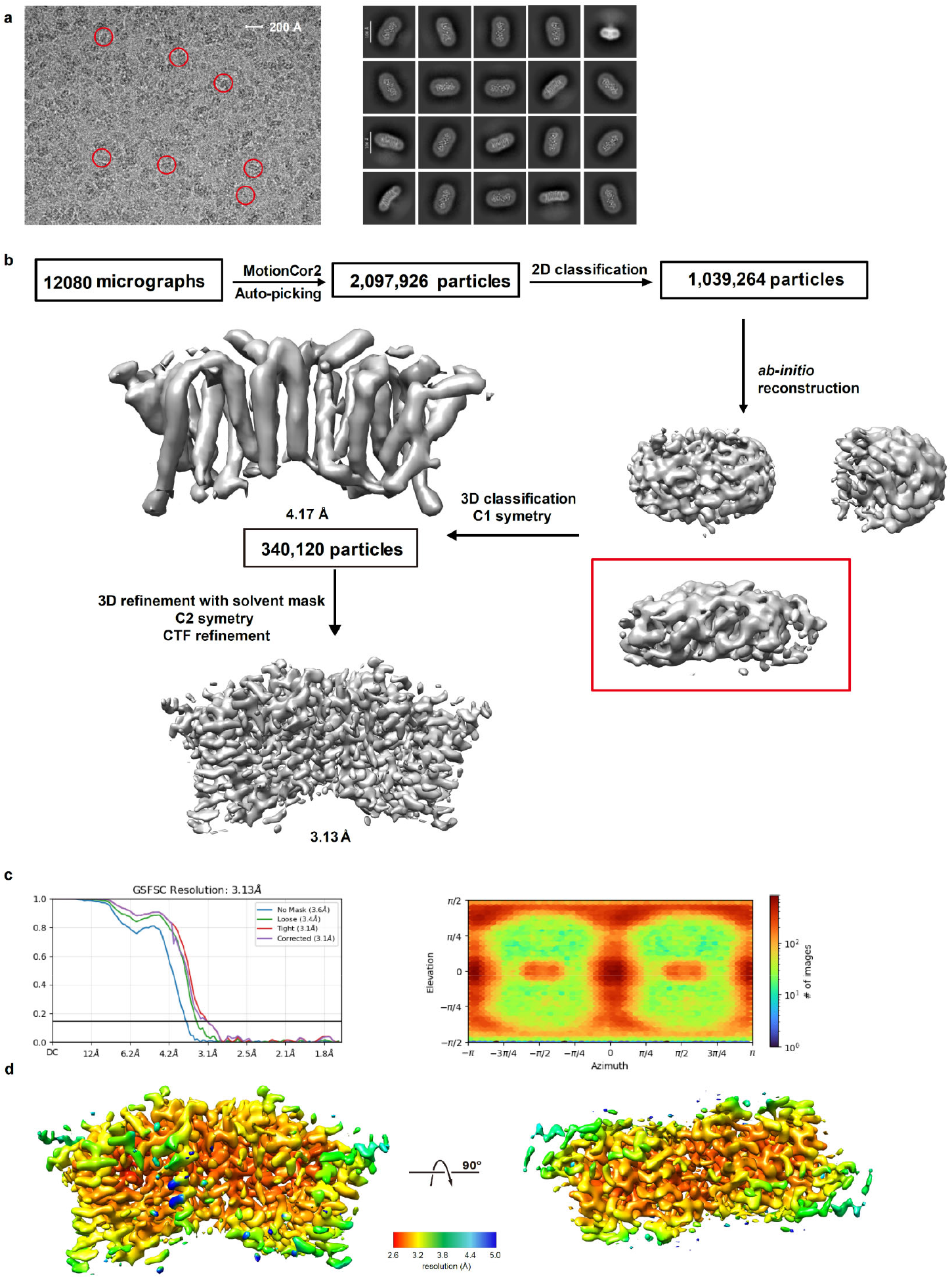
Data processing and map resolution of hPTDSS1 data set. **a**. A representative micrograph of ecPTDSS2 (left) and representative 2D class averages (right). Representative particles are highlighted in red circles. **b**. A flowchart for data processing and the final map. **c**. The gold-standard Fourier shell correlation curve for the final map. **d**. Local resolution map shown in two orientations.

**Extended Data Figure 8.**
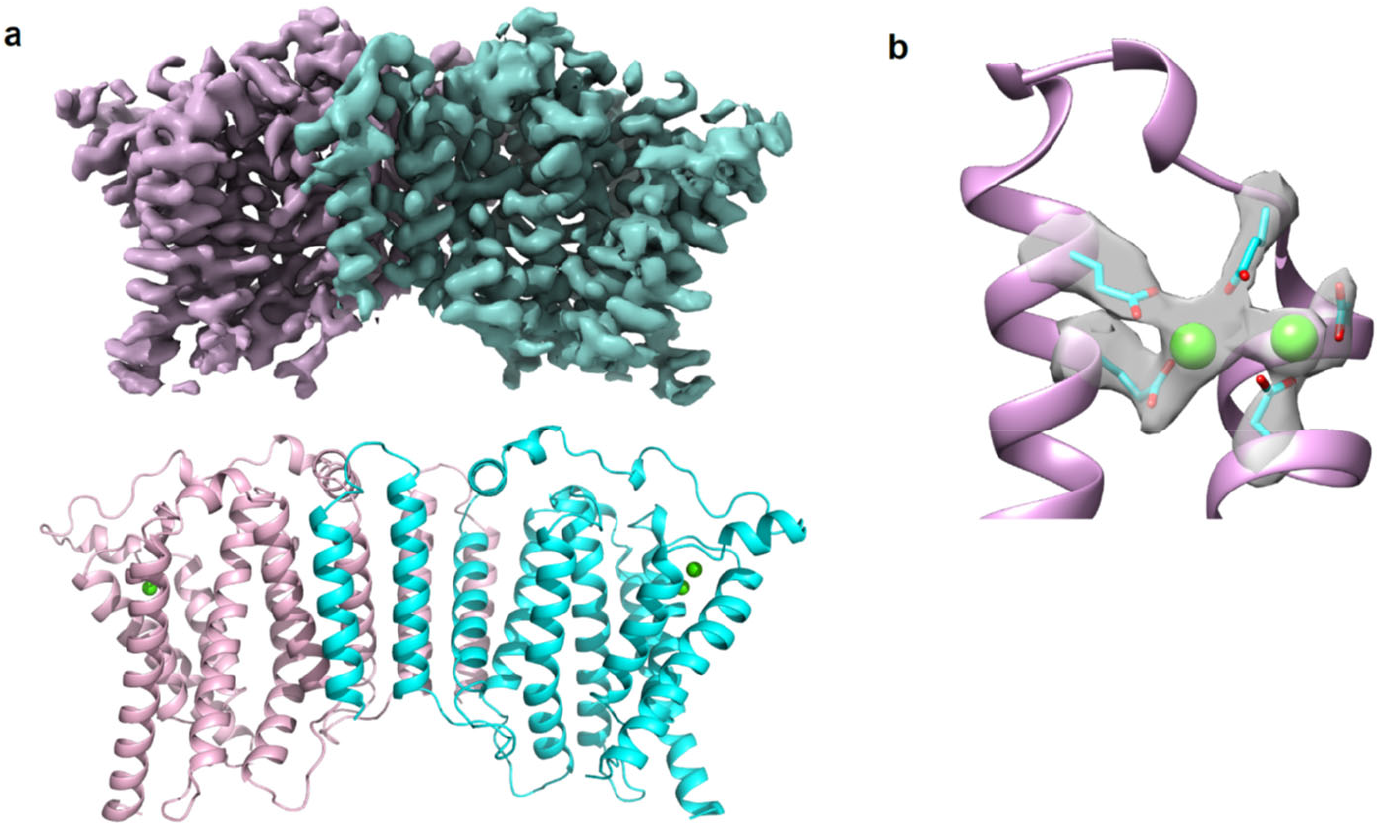
Structure of hPTDSS1 and Ca site. **a**. Density map and model of human PTDSS1 structure determined in this study. **b**. Density of Ca^2+^ site in hPTDSS1. Protein is shown in carton, residues coordinating Ca^2+^ are shown as cyan sticks, Ca^2+^ ions are shown as green spheres. Density is shown in transparent gray surface.

**Extended Data Figure 9.**
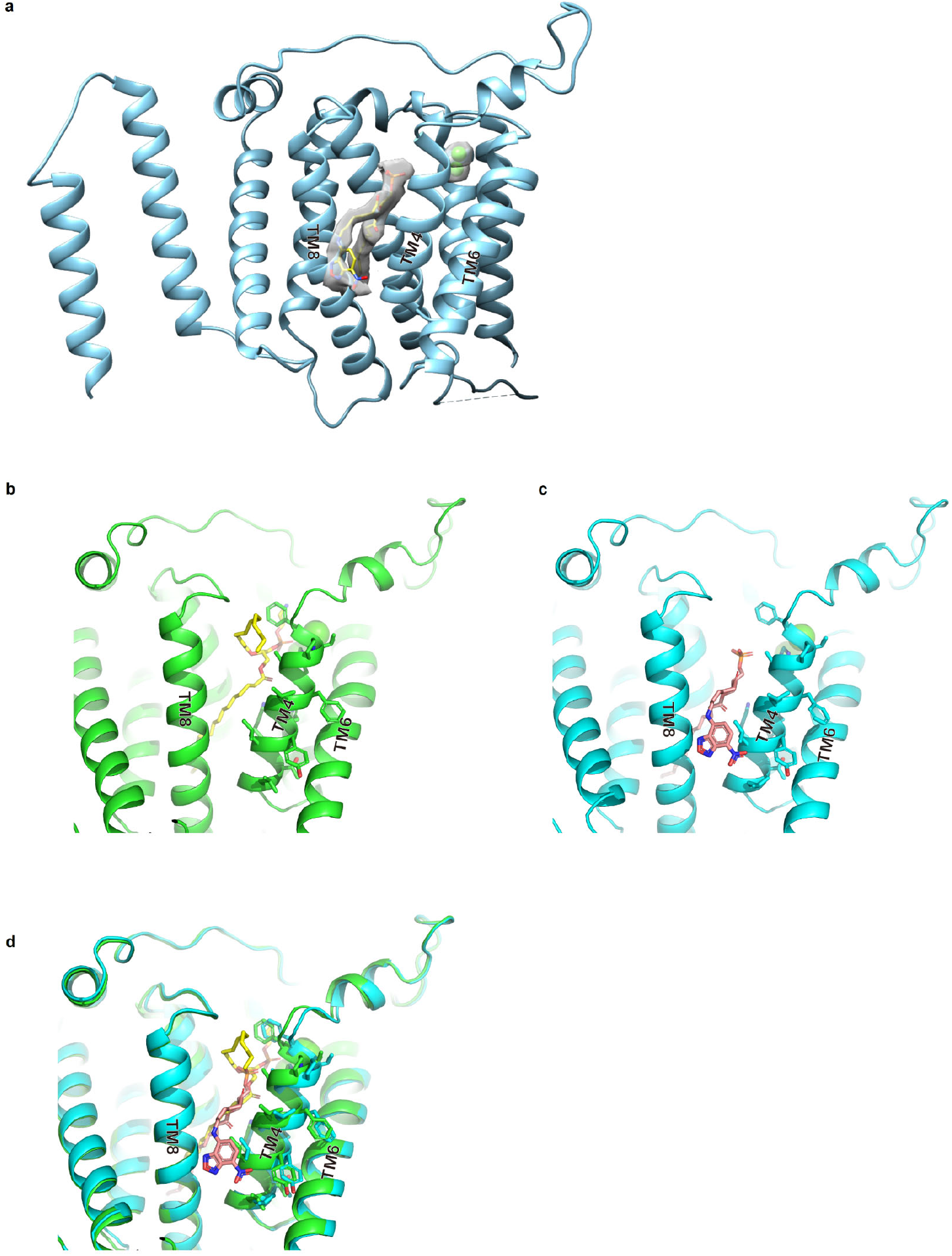
Structural comparison between ecPTDSS2-POPE and ecPTDSS2-NBD-PA. **a**. Structure of ecPTDSS2-NBD-PA is shown as cyan cartoon, with NBD-PA shown as yellow sticks and Ca^2+^ atom as green sphere, and their density shown as gray surface. **b-c**. Structures comparison side by side. Structure of ecPTDSS2 with POPE is shown in **b**, with POPE shown as yellow sticks. Structure of ecPTDSS2-NBD-PA is in **c**, with NBD-PA shown as pink sticks. **d**. Alignment of ecPTDSS2-PE and ecPTDSS2-NBD-PA structure, with obvious shift on TM4.

**Extended Data Figure 10.**
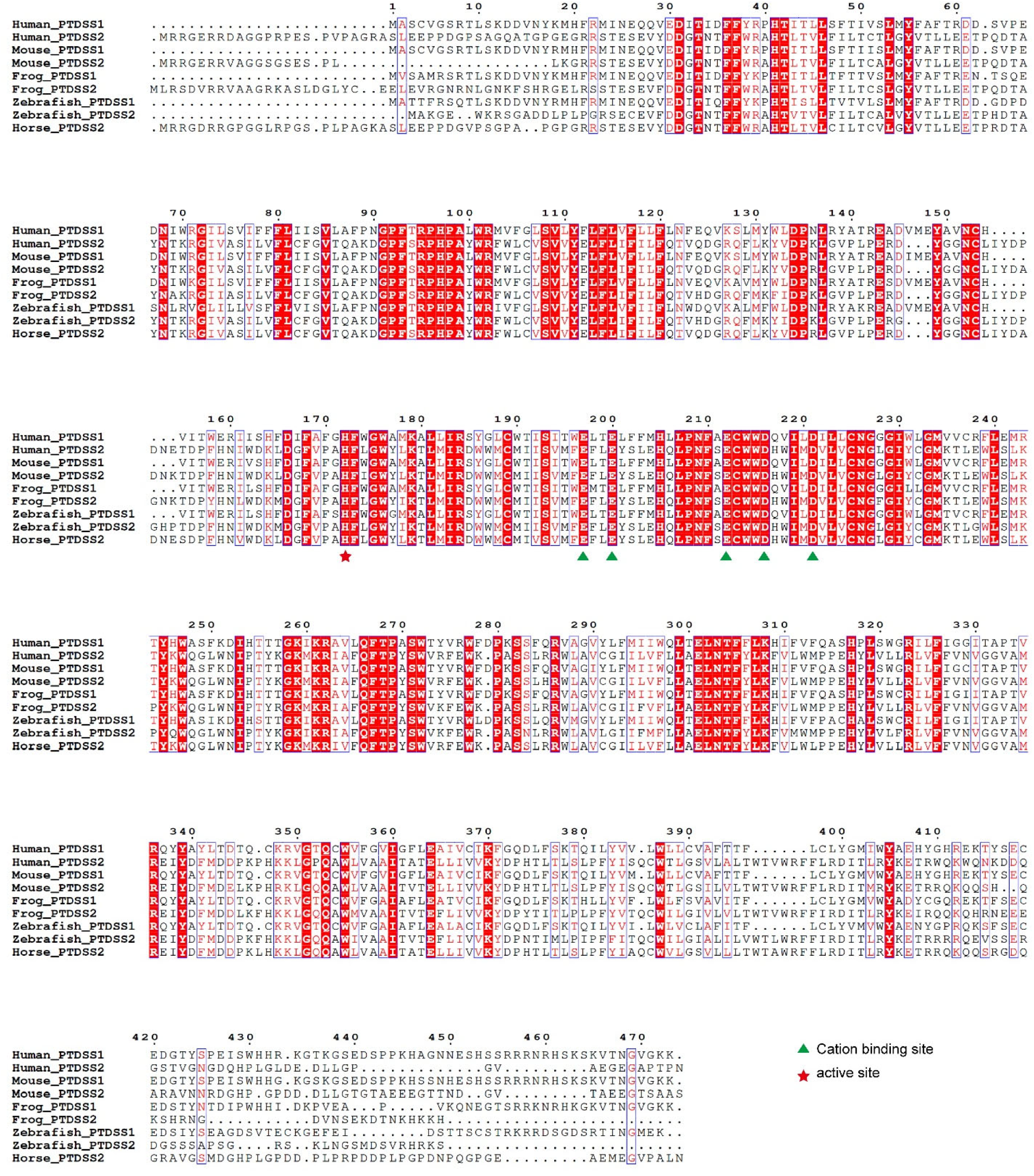
Sequence alignment. Human PTDSS1(P48651 [https://www.uniprot.org/uniprotkb/P48651/entry]), human PTDSS2 (Q9BVG9 [https://www.uniprot.org/uniprotkb/Q9BVG9/entry]), mouse PTDSS1 (Q99LH2 [https://www.uniprot.org/uniprotkb/Q99LH2/entry]), mouse PTDSS2 (Q9Z1X2 [https://www.uniprot.org/uniprotkb/Q9Z1X2/entry]), frog PTDSS1 (B1H3H9 [https://www.uniprot.org/uniprotkb/B1H3H9/entry]), frog PTDSS2 (B5DE41 [https://www.uniprot.org/uniprotkb/B5DE41/entry]), zebrafish PTDSS1 (Q803C9 [https://www.uniprot.org/uniprotkb/Q803C9/entry]), zebrafish PTDSS2 (E7EY42 [https://www.uniprot.org/uniprotkb/E7EY42/entry]), horse PTDSS2 (F6YG08 [https://www.uniprot.org/uniprotkb/F6YG08/entry]) are aligned using the Clustal Omega server^34^. Secondary structural elements of xlCHPT11 are marked above the alignment. Residues are colored based on their conservation using the ESPript server^35^. Cation binding site residues are marked in green triangles. Residue in the active site are marked with red stars.

**Extended data table 1.** Summary of cryo-EM data collection, processing, and structural refinement.

|  | ecPTDSS2-Serine<br>(EMD-75056)<br>(PDB 10BY) | ecPTDSS2-POPE<br>(EMD-75054)<br>(PDB 10BW) | ecPTDSS2-NBD-PA<br>(EMD-75055)<br>(PDB 10BX) | hPTDSS1<br>(EMD-75057)<br>(PDB 10BZ) |
| --- | --- | --- | --- | --- |
| **Data collection and processing** |  |  |  |  |
| Magnification | 105,000 | 105,000 | 105,000 | 105,000 |
| Voltage (kV) | 300 | 300 | 300 | 300 |
| Electron exposure (e-/Å<sup>2</sup>) | 50 | 50 | 50 | 50 |
| Defocus range (μm) | [-2.0, -0.8] | [-2.0, -0.8] | [-2.0, -0.8] | [-2.0, -0.8] |
| Pixel size (Å) | 1.06 | 1.07 | 0.832 | 1.07 |
| Symmetry imposed | C2 | C2 | C2 | C2 |
| Initial particle images (no.) | 8,734,375 | 6,266,551 | 6,602,250 | 2,097,926 |
| Final particle images (no.) | 347,969 | 279,188 | 482,496 | 340,120 |
| Map resolution (Å) | 2.9 | 3.2 | 2.8 | 3.1 |
| FSC threshold | 0.143 | 0.143 | 0.143 | 0.143 |
| Map resolution range (Å) | 2.4-4.0 | 2.7-3.5 | 2.4-4.4 | 2.6-5.0 |
| **Refinement** |  |  |  |  |
| Initial model used (PDB code) | AlphaFold2 | 10BY | 10BY | AlphaFold2 |
| Model resolution (Å) | 3.3 | 3.5 | 3.3 | 3.6 |
| FSC threshold | 0.5 | 0.5 | 0.5 | 0.5 |
| Model resolution range (Å) | 2.5-3.2 | 2.7-3.4 | 2.5-3.3 | 2.9-3.4 |
| Map sharpening *B* factor (Å<sup>2</sup>) | -110 | -130 | -100 | -130 |
| Model composition |  |  |  |  |
| Non-hydrogen atoms | 6166 | 6480 | 6418 | 5670 |
| Protein residues | 700 | 700 | 712 | 678 |
| Ligands | 14 | 24 | 20 | 4 |
| *B* factors (Å<sup>2</sup>) |  |  |  |  |
| Protein | 69 | 56.8 | 119 | 117 |
| Ligand | 12.7 | 35.1 | 50.1 | 96 |
| R.m.s. deviations |  |  |  |  |
| Bond lengths (Å) | 0.003 | 0.003 | 0.011 | 0.003 |
| Bond angles (°) | 0.625 | 0.551 | 0.910 | 0.547 |
| Validation |  |  |  |  |
| MolProbity score | 2.01 | 1.62 | 1.65 | 1.85 |
| Clashscore | 7.38 | 4.89 | 4.83 | 7.37 |
| Poor rotamers (%) | 1.9 | 0 | 0 | 0 |
| Ramachandran plot |  |  |  |  |
| Favored (%) | 94.2 | 94.8 | 94.0 | 93.1 |
| Allowed (%) | 5.2 | 4.4 | 6.0 | 6.5 |
| Disallowed (%) | 0.6 | 0.8 | 0 | 0.4 |

